# Postnatal developmental temperature affects the ontogeny of heat- and cold tolerance

**DOI:** 10.1101/2025.11.07.686921

**Authors:** Elin Persson, Maria Correia, Andreas Nord

## Abstract

Thermoregulatory precision is key to maintaining fitness in thermally unstable environments. Previous research found that development in the warmth renders birds better able to handle mild heat stress, while cold postnatal development gives no apparent benefit upon mild cold exposure. It is still not known how developmental temperature affects maximal heat tolerance limits, or if there is a physiological trade-off between heat and cold tolerance. We investigated how postnatal developmental under simulated cold snap- or heatwave-like conditions from hatching until reproductive maturity impacted the ontogeny of maximal heat- and cold tolerance in Japanese quail (*Coturnix japonica*). To study if any such effects were reversible or permanently programmed, we transferred half of each treatment group to intermediate common garden conditions once reproductive maturity was reached and repeated the measurements several weeks later. Development in heatwave-like conditions increased evaporative water loss rate and moved heat tolerance limits upwards, whereas cold snap-like development rendered more thermogenic birds with improved cold tolerance limits. However, we found no evidence for a trade-off between thermogenic performance and evaporative cooling capacity. The common garden birds converged in nearly all thermoregulatory traits at the end of the study, suggesting that the prior emergence of temperature-dependent phenotypes reflected reversible plasticity. We suggest that improved temperature tolerance limits improve performance in matched thermal conditions, reducing the rate at which thermal injury accrues. Yet, in the short term, we found markedly lower capacity to acclimate heat tolerance compared to cold tolerance, with possible implications for life in a warming world.

## Introduction

Accumulating evidence suggests that both unusually high or low temperature impact nearly every aspect of endotherm performance, ranging from offspring growth, maturation and survival (Cunningham et al., 2013; Nord & Nilsson, 2011; Salaberria et al., 2013) to adult behaviour and reproduction (Cunningham et al., 2021; Edwards et al., 2015; Nilsson & Nord, 2018; Nord & Nilsson, 2019). It follows that improved thermoregulatory precision is key to maintaining fitness in thermally labile environments. In ostriches (*Struthio camelus*), for example, a more pronounced capacity for heat dissipation leads to higher egg laying rates during hotter conditions compared to less capable heat dissipating females (Svensson et al., 2023). Similarly, black-capped chickadees (*Poecile atricapillus*) with higher maximal thermogenic capacity had higher probability of overwinter survival compared with less thermogenic birds (Petit et al., 2017).

The importance of body temperature (*T*_b_) control in a changing world raises the question of the ontogeny of thermoregulatory phenotypes. It has been hypothesised that thermal conditions during the developmental period prime subsequent temperature tolerance through permanent modification of key physiological pathways (e.g., hormonal control of thermogenesis and homeostasis [Bize et al., 2010; Piestun et al., 2008; reviewed by Ruuskanen et al., 2021]). This has often been interpreted as thermal adaptation that increases fitness when the developmental and post-developmental conditions overlap (reviewed by Nichelmann, 2004; Nord & Giroud, 2020), but carries potential costs when these environments are mismatched (Nord & Giroud, 2020). By comparison, the long-term consequences of peri- and postnatal temperature on thermoregulatory traits are not well understood. Research has shown that Japanese quail (*Coturnix japonica*) reared in heatwave-like temperatures were better at handling mild heat stressors, specifically by reducing metabolic heat production (MHP) and elevating evaporative cooling capacity (ECC), as juveniles (Persson et al., 2024). Analogous exposure to cold snap-like postnatal temperature did not result in any thermoregulatory changes upon exposure to mild cold. Nevertheless, measurements below thermal limits may not necessarily capture the full dynamics of how developmental temperature influences heat- and cold tolerance.

Almost all avian heat tolerance studies have been performed in hot and dry climates, on the assumption that animals in such biomes have low capacity to accommodate additional thermal cstress. However, even high-latitude birds overheat frequently (Nilsson & Nord, 2018) and experimental relief from this risk increases fitness by re-investment of resources otherwise spent on thermoregulation into both current and future reproduction (Andreasson et al., 2020; Nord & Nilsson, 2019). The fitness costs of warming may be even more severe at high latitude, where animals are poorly equipped to deal with heat stress (Nord & Folkow, 2018; O’Connor et al., 2021), possibly on account of weak historic selection for high temperature tolerance. In line with this, recent work shows that, in ostriches, there is a negative genetic correlation between heat- and cold tolerance at the level of reproductive effort (Schou et al., 2022) that may be further modified by local adaptation of thermoregulation experienced in the natural ranges of different subspecies (Svensson et al., 2023). In contrast, studies on the fruit fly (*Drosophila melanogaster*) show that there is a positive correlation between heat and cold tolerance (Bubliy & Loeschcke, 2005). This is less likely to occur in endotherms, because acclimation to high temperature typically involves downregulation of metabolic rate to lessen heat gain (Collin et al., 2001; Wojciechowski et al., 2021) whereas cold-acclimation is associated with upregulated heat production rate to support thermogenesis. To the best of our knowledge, no endotherm study has yet addressed whether the purported trade-off between heat and cold limits has a physiological basis.

We investigated whether developmental temperatures impact the ontogeny of maximal heat- and cold tolerance limits. Japanese quail was reared in cold snap (10°C) or heatwave-like (30°C) temperature conditions from hatching until reproductive maturity, after which half of each treatment group were transferred to common garden conditions at intermediate temperature (20°C) to test whether any temperature-dependent phenotypes stemmed from permanent programming or reversible plasticity. Cold- and heat tolerance were measured halfway to asymptotic size during the exponential growth phase, around the start of reproductive maturity, and again well into adulthood. We predicted that birds growing up under warm conditions would cope better in high ambient temperatures, evaporating water at a higher rate and having a higher heat tolerance limit, compared to birds from cold conditions. By contrast, we predicted that cold-rearing would increase thermogenic capacity and improve the cold tolerance limit relative to warm-rearing. If these changes were caused by permanent programming of the thermoregulatory phenotype, we predicted that trait values of the common-garden birds would reflect rearing-, but not housing-, temperature. However, if there was no programming, we predicted that the effects would return to pre-acclimation values in the common garden. By measuring empirically the relationship between heat- and cold-tolerance at both the population- and individual levels, our design also permitted testing for the first time whether the recently reported reproductive trade-off between these traits (Schou et al., 2022) has a thermoregulatory basis.

## Methods

### Species and husbandry

Japanese quail eggs (n=85) were bought from a commercial breeder (Sigvard Månsgård, Åstorp, Sweden), and placed in an automatic turner that turned the eggs for 5 min every 1 h at room temperature (18-20°C) for 1-7 days. Then, the eggs were put into one of two Brinsea Ova Easy 190 incubators (6 – 8 eggs per day; Brinsea, Weston-super-Mare, United Kingdom) at 37.45°C ± 0.358 or 37.55°C ± 0.205 (mean ± s.d.) and 50% relative humidity (automatically regulated using a Brinsea Ova-Easy Advance Humidity Pump). Eggs were moved from incubation trays to a hatching tray inside the incubator at day 16 of incubation and were checked for hatching twice daily from day 17. Fifty-five eggs of 85 hatched after an incubation period of 19 days (mean ± s.d.: 19.27 ± 0.73 days; range: 18-22 days).

When completely dry (1 to 12 h after hatching), the chicks were banded with uniquely numbered leg bands and randomly allocated to either a cold snap-like (10°C; 9.70 ± 0.31°C [mean ± s.d.]; n=25; Table S1) or a heatwave-like housing regime (30°C; 30.02 ± 0.82°C [mean ± s.d.]; n=24; Table S1) in line with previous research (Correia et al., 2025; Persson et al., 2024). The birds remained in these thermal environments until 9 weeks old. Then, half of the of the birds (Cold: n=12, henceforth Cold-mild; Warm: n=11, henceforth Warm-mild) were transferred to a common garden at 20°C (20.01 ± 0.35°C [mean ± s.d.]), whereas the other half remained in their original temperature treatments (Cold: n=11; Warm: n=11) until the end of the experiment (Fig. S1). The birds were housed in open pens (310 × 120 × 60 cm) with food (turkey starter until 3 weeks [Kalkonfoder Start, Lantmännen, Stockholm, Sweden; 25.5% protein], and turkey grower from 3 weeks onwards [Kalkonfoder Tillväxt, Lantmännen, Stockholm, Sweden; 22.5% protein]), water, sand and seashells available *ad libitum*. Mealworms, or lettuce and shredded carrots, were provided as diet supplements daily on alternate days. A heat lamp (35-39°C), compensating for the absence of a brooding female, was available until 2 weeks of age. Body mass (± 0.1 g) and wing length (± 0.5 mm) were measured once weekly and at 1, 2, 3, 8 and 12 weeks of age, respectively. Analyses and results pertaining to morphological data are disclosed in the ESM.

### Measurement of body temperature

When the birds were 13 to 16 days old, we implanted a sterile temperature-sensitive passive integrated transponder (< 0.5% of body weight; LifeChip BioTherm, Destron Fearing, South St. Paul, MN, USA) into the intraperitoneal cavity following previously described procedures (Persson et al., 2024). This permitted non-contact measurement of *T*_b_ during cold- and heat tolerance measurements (see below). *T*_b_ was also measured in the holding pens in-between experimental periods (at 5-7 and 10-12 weeks). Since there were, at most, minor differences between treatment groups, these data are not considered further below (see ESM for details).

### Respirometry

Respirometry measurements to derive cold- and heat tolerance were performed during daytime between 29 March and 20 June 2022. Representative raw data for these experiments are presented in Fig. S2.

We defined maximum thermogenic performance (summit metabolic rate, *M*_sum_) as the highest metabolic rate reached during sliding cold exposure, and the cold tolerance limit as the temperature at which *M*_sum_ was attained. These measurements were taken at 3, 8 and 12 weeks (Fig. S1; Table S1). A 13 L glass respirometry chamber contained inside a climate test chamber (Weiss Umwelttechnik C180, Reiskirchen, Germany) was ventilated with a dry gas mixture of 79% helium and 21% oxygen (‘helox’; 4.44 ± 0.03 L/min [mean ± s.e.m.], measured using an Alicat 20SLPM mass flow meter [Alicat Scientific Inc., Tucson, AZ, USA]) from which we subsampled 128.27 ± 1.25 ml/min (mean ± s.e.m.; SS4 pump, Sable Systems, Las Vegas, NV, USA) to measure water vapour density and oxygen consumption (using RH300 and FC-10 analysers, respectively; Sable Systems). Water vapour and carbon dioxide were removed from the airstream before measuring oxygen consumption, using drierite and ascarite, respectively. After thermal acclimation (15 min; 3 weeks at 5°C; 8 and 12 weeks at −5°C), chamber temperature was acutely decreased to −5°C (3 weeks) or −10°C (8 and 12 weeks) after which temperature decreased continuously by 20°C per hour. An experiment ended; a) when oxygen consumption started to decrease for a decrease in chamber temperature; or b) when there was no change in oxygen consumption for at least 20 min whilst ambient temperature decreased (Fig. S2A). Immediately after, the bird was placed in a small cage (52 × 32 × 30 cm) with access to water and a heat lamp where they remained until being transferred back to their pen 20 to 40 min later.

We defined the heat tolerance limit as the temperature above which the bird could no longer increase its evaporative water loss (EWL) in response to heating, and derived maximal evaporative capacity based on EWL recorded at this temperature. Heat tolerance measurements were taken 4-9 days after the cold tolerance assay starting at 4, 9 and 13 weeks (Fig. S1; Table S1). The same instruments were used for both assays. For heat tolerance, dry (drierite) atmospheric air (10.06 ± 0.01 L/min [mean ± s.e.m.]) was pushed through the 13 L chambers of which 403.33 ± 0.28 ml/min (mean ± s.e.m.) was subsampled for analyses of water vapor density and oxygen. A metal grid platform was placed inside the respirometry chamber over a 1.5 – 2 cm layer of mineral oil, to remove any influence of evaporation from faeces on EWL. After acclimation in 30°C (for 30 to 60 min until stable gas traces), chamber temperature was acutely raised to 40°C, after which it was increased in 2°C increments as soon as gas traces had remained stable for at least 5 min (Noakes et al., 2016; Talbot et al., 2018). Experiments ended: a)when oxygen consumption or EWL decreased sharply when chamber temperature increased; b)if *T*_b_ increased > 45°C; or c)if birds showed signs of stress or loss of coordination (McKechnie et al., 2016; Whitfield et al., 2015; Fig S2B). Immediately after, EtOH (70%) was sprayed onto the ventral plumage and skin and the birds were placed in front of a fan (Whitfield et al., 2015) in the small cage described above, with water *ad libitum* until being transferred to the holding pen 20 to 40 min later. Birds had recovered *T*_b_ within minutes of being placed in front of the fan (Author, pers. obs.).

During all tolerance assays, we measured air temperature inside the chambers at floor and ceiling level using copper-constantan thermocouples (36-gauge type T) attached to a TC-2000 thermocouple box (Sable Systems) and manually adjusted the climate chamber settings to keep bird temperature at target. All respirometry sessions started and ended with a recording of reference air (i.e., baseline).

### Data handling

Of the 55 birds that hatched, 6 died within the first week of life, and another 5 birds died of natural causes during the experiment. Two birds were never considered for the metabolic measurements since additional measurements could not be fitted within the birds’ photophase. One bird was removed prematurely from the heat tolerance measurements at 4 weeks due to stress and was not considered further in the analyses. At 12 weeks, one bird was removed from wing length analyses because its primary feathers were damaged. Final sample sizes for each measurement period and trait are presented in Table S1.

Respirometry data were extracted using ExpeData (v. 1.9.27; Sable Systems). In cold tolerance measurements, we extracted mean values of the 10 min preceding the endpoint (Swanson et al., 1996). In heat tolerance measurements, we extracted data corresponding to the most stable 2 min of oxygen consumption in the final measurement temperature. Oxygen consumption (ml min^-1^) was calculated using eq. 11.1 in Lighton (2008) and was converted to MHP (W) by assuming an energy equivalence of 20 J per 1 mL of O_2_ (Kleiber, 1961). EWL (g h^-1^) was calculated using eq. 11.9 in Lighton (2008) and converted to evaporative heat loss (EHL; W) by assuming that evaporation of 1 ml water requires 2406 J (Wallace & Hobbs, 2006). ECC was calculated as the ratio between EHL and MHP (Lasiewski et al., 1966). This metric strongly predicted the heat tolerance limit at all ages (Fig. S3; Table S2).

### Statistics

All statistical analyses were performed using R ver. 4.3.0 (R Core Team, 2023). Linear mixed models (lmer() in lme4; Bates et al., 2015) were used to analyse whether thermal and metabolic responses differed between the cold and warm treatment groups. MHP, *M*_sum_, EWL, ECC, heat- and cold tolerance limits and *T*_b_ were used as response variables, with treatment, age, and treatment×age as factors. Body mass (mean centred by treatment and age) was used as a continuous covariate. Bird ID was used as a random intercept to account for repeated measurements. The effects of the transfer to common garden were analysed at the final metabolic measurement period using ANOVA (anova() function in stats package), comparing Warm-mild birds against Warm birds, and Cold-mild birds against Cold birds. All dependent variables but tolerance limits and *T*_b_ were log-transformed in all analyses above to meet parametric assumptions.

The phenotypic association between ECC and heat-producing capacity (i.e., *M*_sum_), potentially indicative of a trade-off between heat- and cold tolerance, was addressed using Pearson correlation (cor.test() in the stats package). Separate correlations were applied to each age interval, i.e., 3-4, 8-9, and 12-13 weeks of age.

*P*-values from linear mixed models were assessed using likelihood ratio tests. Non-significant interactions were removed from the models, but all main effects were kept. Significant effects were compared using a post hoc test (pairs() in the emmeans package; Lenth, 2023) and estimates were back-transformed when applicable. Significant effects from ANOVAs were compared using a Tukey post hoc test (glht() in the multcomp package; Hothorn et al., 2008).

## Results

### Cold tolerance

Birds that grew up under cold conditions had significantly higher log(*M*_sum_) at 8 (17.2%) and 12 (13.0%) weeks of age than birds from warm conditions, but this effect was absent at 3 weeks (Treatment×Age interaction: p = 0.059; Fig. 1A; Table 1). When the interaction was removed from the model, Cold birds had 10.9% higher log(*M*_sum_) than Warm birds (Fig. 1A; Table 1). In addition, log(*M*_sum_) increased with age and was 37.8 % higher at 8 weeks and 43.5% higher at 12 weeks than at 3 weeks of age (Fig. 1A; Table 1). The cold tolerance limit was affected by the interaction between treatment and age. Specifically, Cold birds reached significantly lower temperature than Warm birds at 8 (by 5.4°C) and 12 (by 6.9°C) weeks of age, but there was no significant difference at 3 weeks (Fig. 1B; Table 1). *T*_b_ at the cold tolerance limit was not affected by the interaction between treatment and age nor by treatment or age alone (Fig. S4A; Table 1).

**Table 1.** Parameter estimates explaining the effects of developing in either Cold (10°C) or Warm (30°C) temperature conditions in Japanese quail. Estimates, test statistics, degrees of freedom, P-values and standard deviations (σ) for the random factor from linear mixed models at 3 different ages, in cold-(measured in helox) and heat tolerance measurements. Back-transformed model estimates for log-transformed variables are shown in the table. Statistics for main effects is not provided when the interaction between treatment and age was significant. Significant (P < 0.05) post hoc comparisons are shown by different letters within brackets. Abbreviations: *M*_sum_: Summit metabolic rate, MHP: Metabolic heat production; EWL: Evaporative water loss; ECC: Evaporative cooling capacity; *T*_b_: Body temperature.

| Model | Estimate ± s.e.m. | LR | Df | P | $\sigma_{ID}$ \| $\sigma_{total}$ |
| --- | --- | --- | --- | --- | --- |
| **Cold tolerance** |  |  |  |  |  |
| <u>Log($M_{sum}$)</u> |  |  |  |  |  |
| Treatment:Age |  | 5.66 | 2 | 0.0591 . |  |
| 3 weeks |  |  |  |  |  |
| Cold [A] | 6.80 ± 0.150 | | | | |
| Warm [A] | 6.46 ± 0.152 | | | | |
| 8 weeks |  |  |  |  |  |
| Cold [A] | 9.86 ± 0.250 | | | | |
| Warm [B] | 8.41 ± 0.225 | | | | |
| 12 weeks |  |  |  |  |  |
| Cold [A] | 10.10 ± 0.354 | | | | |
| Warm [B] | 8.94 ± 0.299 | | | | |
| Treatment |  | 15.92 | 1 | <0.0001 *** |  |
| Cold [A] | 8.75 ± 0.144 | | | | |
| Warm [B] | 7.89 ± 0.134 | | | | |
| Age |  | 126.77 | 2 | <0.0001 *** |  |
| 3 weeks [A] | 6.62 ± 0.108 | | | | |
| 8 weeks [B] | 9.12 ± 0.170 | | | | |
| 12 weeks [B] | 9.50 ± 0.234 | | | | |
| Body mass | 1.003 ± 1.001 | 17.94 | 1 | <0.0001 *** | |
| Bird ID (random) |  | 0.32 | 1 | 0.5723 | 1.00 \| 2.01 |
| <u>Cold tolerance limit (°C)</u> |  |  |  |  |  |
| Treatment:Age |  | 6.95 | 2 | 0.0310 * |  |
| 3 weeks |  |  |  |  |  |
| Cold [A] | -11.18 ± 0.897 | | | | |
| Warm [A] | -9.43 ± 0.957 | | | | |
| 8 weeks |  |  |  |  |  |
| Cold [A] | -26.78 ± 1.058 | | | | |
| Warm [B] | -21.43 ± 1.088 | | | | |
| 12 weeks |  |  |  |  |  |
| Cold [A] | -29.80 ± 1.417 | | | | |
| Warm [B] | -22.94 ± 1.351 | | | | |
| Body mass |  | 3.30 | 1 | 0.0692 |  |
| Bird ID (random) |  | 1.48 | 1 | 0.2231 | 3.42 \| 20.11 |
| <u>$T_b$ (°C)</u> |  |  |  |  |  |
| Treatment |  | 0.35 | 1 | 0.5517 |  |
| Age |  | 0.40 | 2 | 0.8200 |  |
| Treatment:Age |  | 1.81 | 2 | 0.4048 |  |
| Body mass |  | 0.003 | 1 | 0.9551 |  |
| Bird ID (random) |  | 8.86 | 1 | 0.0029 ** | 0.62 \| 1.67 |
| **Heat tolerance** |  |  |  |  |  |
| <u>Log(MHP)</u> |  |  |  |  |  |
| Treatment:Age |  | 6.55 | 2 | 0.0379 * |  |
| 4 weeks |  |  |  |  |  |
| Cold [A] | 3.21 ± 0.140 |  |  |  |  |
| Warm [A] | 2.89 ± 0.124 |  |  |  |  |
| 9 weeks |  |  |  |  |  |
| Cold [A] | 3.72 ± 0.159 |  |  |  |  |
| Warm [A] | 3.89 ± 0.170 |  |  |  |  |
| 13 weeks |  |  |  |  |  |
| Cold [A] | 3.66 ± 0.216 |  |  |  |  |
| Warm [A] | 3.97 ± 0.226 |  |  |  |  |
| Body mass | 1.004 ± 1.001 | 15.50 | 1 | <0.0001 *** |  |
| Bird ID (random) |  | 10.47 | 1 | 0.0012 ** | 1.02\|2.04 |
| <u>Log(EWL)</u> |  |  |  |  |  |
| Treatment |  | 7.68 | 1 | 0.0056 ** |  |
| Cold [A] | 3.52 ± 0.113 |  |  |  |  |
| Warm [B] | 4.00 ± 0.130 |  |  |  |  |
| Age |  | 69.71 | 2 | <0.0001 *** |  |
| 4 weeks [A] | 3.15 ± 0.082 |  |  |  |  |
| 9 weeks [B] | 4.02 ± 0.104 |  |  |  |  |
| 13 weeks [B] | 4.17 ± 0.136 |  |  |  |  |
| Treatment:Age |  | 2.96 | 2 | 0.2278 |  |
| Body mass |  | 1.58 | 1 | 0.2094 |  |
| Bird ID (random) |  | 32.62 | 1 | <0.0001 *** | 1.02\|2.03 |
| <u>Log(ECC)</u> |  |  |  |  |  |
| Treatment |  | 3.37 | 1 | 0.0663 |  |
| Age |  | 0.94 | 2 | 0.6258 |  |
| Treatment:Age |  | 3.50 | 2 | 0.1738 |  |
| Body mass | 0.997 ± 1.001 | 4.05 | 1 | 0.0441 * |  |
| Bird ID (random) |  | 34.40 | 1 | <0.0001 *** | 1.05\|2.08 |
| <u>Heat tolerance limit (°C)</u> |  |  |  |  |  |
| Treatment:Age |  | 7.68 | 2 | 0.0215 * |  |
| 4 weeks |  |  |  |  |  |
| Cold [A] | 43.0 ± 0.474 |  |  |  |  |
| Warm [B] | 45.2 ± 0.475 |  |  |  |  |
| 9 weeks |  |  |  |  |  |
| Cold [A] | 44.1 ± 0.467 |  |  |  |  |
| Warm [A] | 44.7 ± 0.481 |  |  |  |  |
| 13 weeks |  |  |  |  |  |
| Cold [A] | 43.5 ± 0.576 |  |  |  |  |
| Warm [B] | 45.3 ± 0.570 |  |  |  |  |
| Body mass |  | 0.77 | 1 | 0.3814 |  |
| Bird ID (random) |  | 36.33 | 1 | <0.0001 *** | 3.51\|4.95 |
| <u>T<sub>b</sub> (°C)</u> |  |  |  |  |  |
| Treatment:Age |  | 0.57 | 2 | 0.0039 ** |  |
| 3 weeks |  |  |  |  |  |
| Cold [A] | 44.65 ± 0.096 |  |  |  |  |
| Warm [B] | 44.41 ± 0.091 |  |  |  |  |
| 8 weeks |  |  |  |  |  |
| Cold [A] | 44.60 ± 0.080 |  |  |  |  |
| Warm [A] | 44.62 ± 0.090 |  |  |  |  |
| 12 weeks |  |  |  |  |  |
| Cold [A] | 44.47 ± 0.111 |  |  |  |  |
| Warm [A] | 44.75 ± 0.114 |  |  |  |  |
| Body mass |  | 11.08 | 1 | 0.4492 |  |
| Bird ID (random) |  | 9.62 | 1 | 0.0019 ** | 0.04\|0.12 |

**Figure 1.**
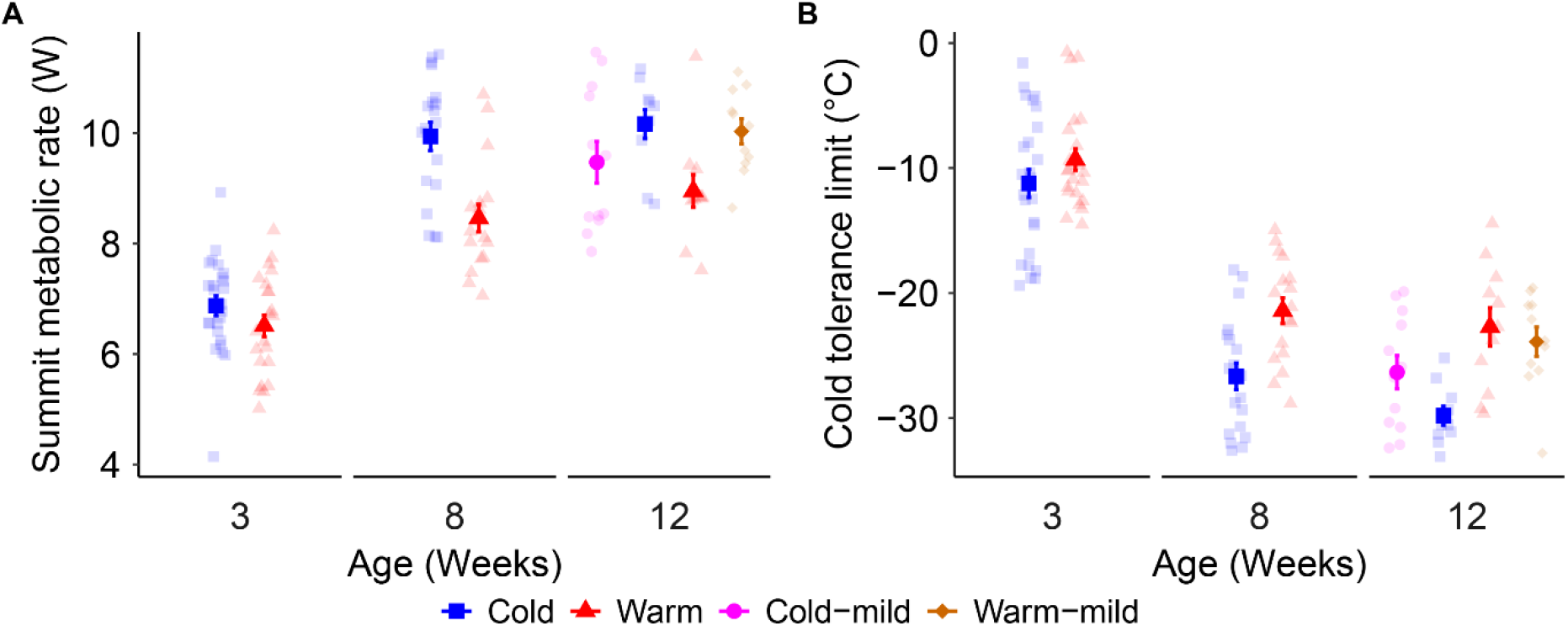
Maximum thermogenic capacity (A) and cold tolerance limit (B) in temperature-acclimated Japanese quail. Birds were raised in either Cold (10°C) or Warm (30°C) conditions until 9 weeks of age. Then, half of each group was transferred to common garden (Cold-mild, Warm-mild; 20°C) and the remainder kept in their original temperature treatments. Sample sizes per age and temperature are stated in Table S1. Points and error bars show estimated means ± s.e.m.. Semi-transparent points show raw data.

### Heat tolerance

There was no significant difference in log(MHP) between Warm and Cold birds at any age. However, log(MHP) was higher at 9 and 13 weeks of age than at 4 weeks. Specifically, Warm birds had significantly higher log(MHP) at 9 and 13 weeks of age (9 weeks: 34.6%; 13 weeks: 37.7%) than at 4 weeks, whereas Cold birds had significantly higher log(MHP) at 9 weeks (15.9%) than at 4 weeks of age (Treatment×Age interaction: p = 0.038). Across ages, Warm-acclimated birds had 13.6% higher log(EWL) than Cold birds (Fig 2B; Table 1). In addition, log(EWL) increased with age, being lower at 4 weeks compared to at the other two ages (Fig. 2B, Table 1). ECC was not affected by the interaction between treatment and age, nor by treatment or age alone. However, there was a tendency (p=0.064) for Warm birds to have higher ECC than Cold birds (Fig. 2C; Table 1). Nevertheless, Warm birds had a higher heat tolerance limit compared to Cold birds (at 4 and 13 weeks [2.2°C and 1.8°C respectively; Fig. 2D, Table 1). *T*_b_ at the heat tolerance limit was affected by the interaction between treatment and age (Fig. S4B; Table 1) such that at 3 weeks, Cold birds had 0.24°C higher *T*_b_ than Warm birds.

**Figure 2.**
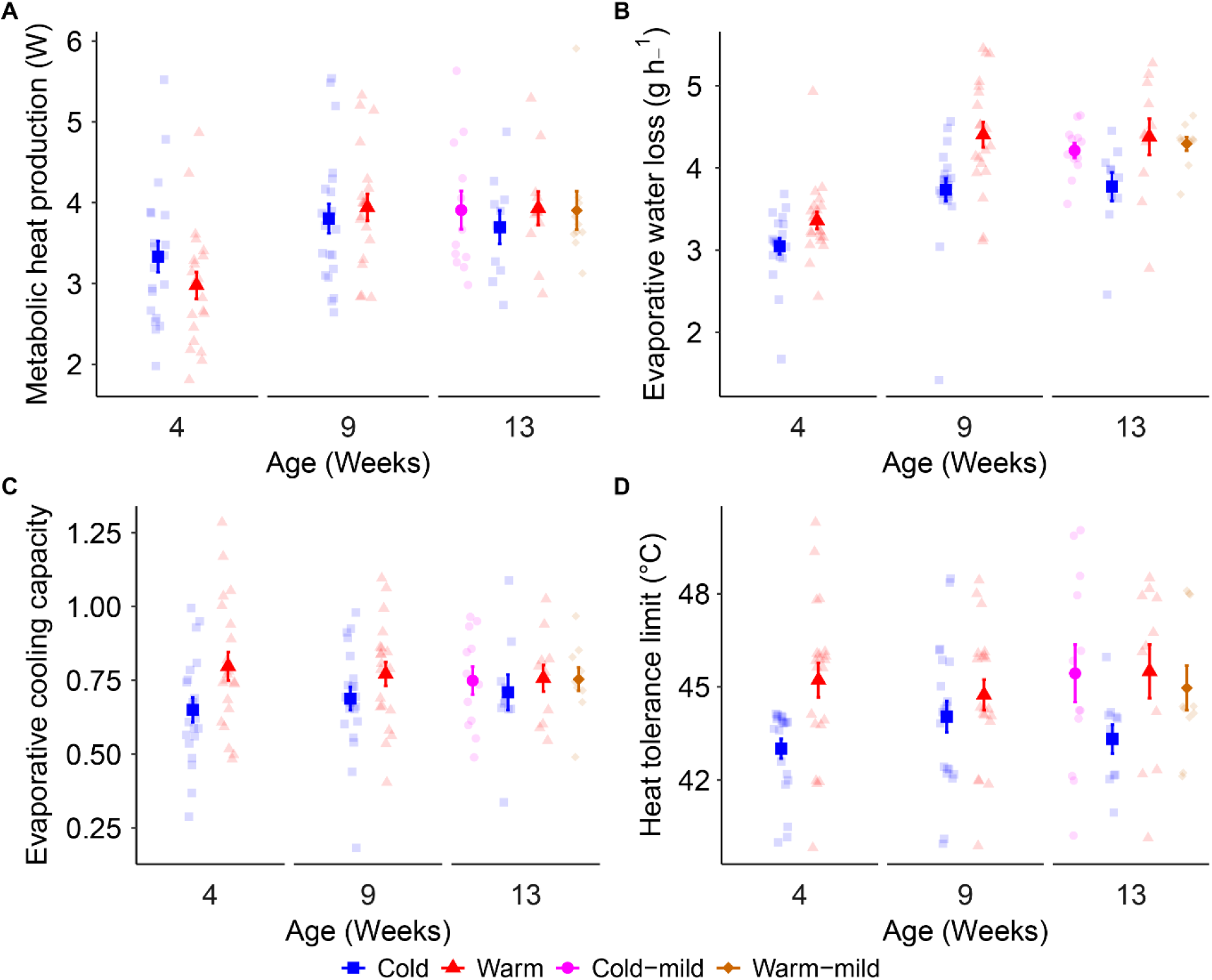
Thermoregulatory traits at the heat tolerance limit in temperature-acclimated Japanese quail. Panels show (A) Metabolic heat production, (B) Evaporative water loss, (C) Evaporative cooling capacity and (D) heat tolerance limit. The birds were raised in Cold (10°C) or Warm (30°C) conditions until 9 weeks of age, after which half of each group were transferred to common garden conditions (Cold-mild, Warm-mild; 20°C). Sample sizes per age and temperature are stated in Table S1. Points and error bars show estimated means ± s.e.m.. Semi-transparent points show raw data.

### Common garden birds

After the transfer to common garden conditions, Warm-mild birds developed 12.4% higher log(*M*_sum_) than birds that remained in Warm conditions (Fig. 1A; Table 2). There was no significant difference in log(*M*_sum_) between Cold-mild and Cold birds. However, Cold-mild birds reduced their cold tolerance limit compared to Cold birds (by 3.5°C; Fig. 1B; Table 2). In contrast, the heat tolerance limit did not differ between Warm and Warm-mild birds. Log(EWL) improved in Cold-mild compared to Cold birds (by 12.6%), with a tendency for an analogous change in the heat tolerance limit (p=0.076; Figs. 2B, E; Table 2). There was no difference in these traits between Warm-mild and Warm birds. *T*_b_ at the tolerance limits did not differ between Cold and Cold-mild birds, nor between Warm and Warm-mild birds (Fig. S4; Table 2).

**Table 2.**
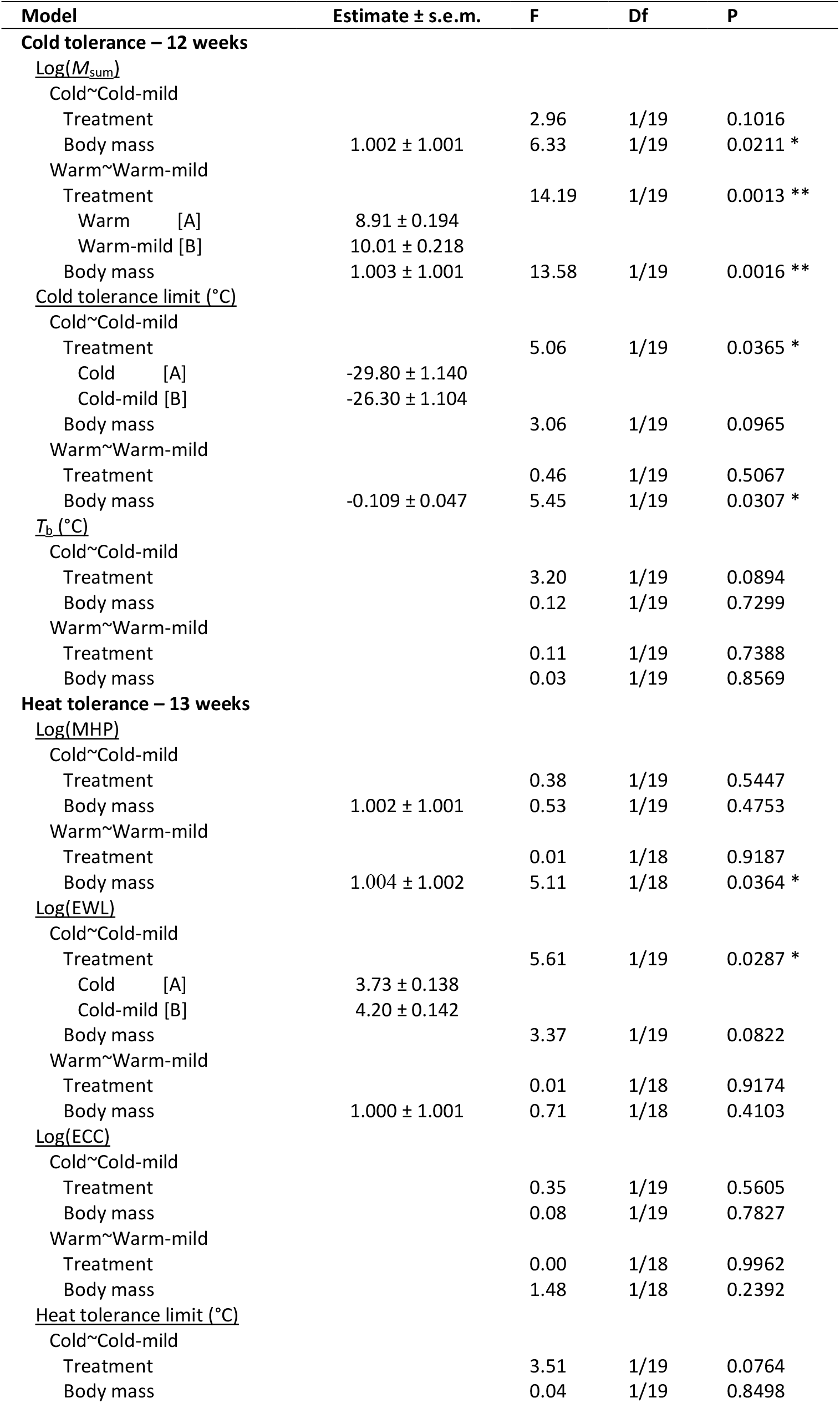
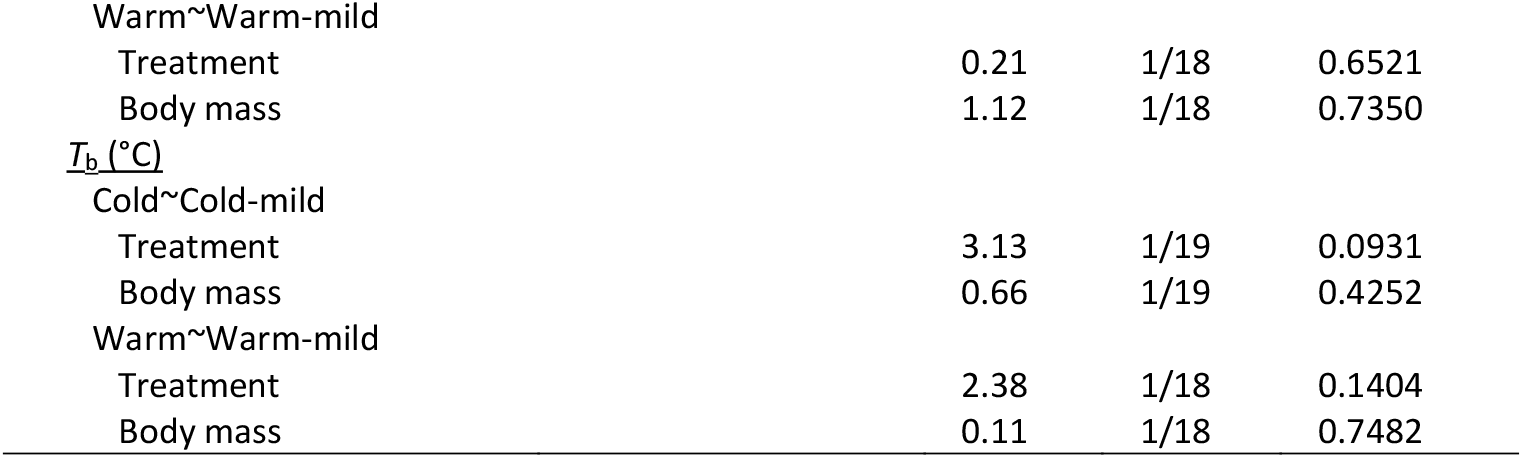
Parameter estimates explaining the effects of moving from either Cold (10°C) or Warm (30°C) treatment to common garden conditions (20°C) in Japanese quail. Estimates, test statistics, degrees of freedom and P-values from ANOVAs at 12 weeks in cold tolerance measurements (measured in helox), and at 13 weeks in heat tolerance measurements. Back-transformed model estimates for log-transformed variables are shown in the table. Significant (P < 0.05) post hoc comparisons are shown by different letters within brackets. Abbreviations: *M*_sum_: Summit metabolic rate; MHP: Metabolic heat production; EWL: Evaporative water loss; ECC: Evaporative cooling capacity; *T*_b_: Body temperature.

### The relationship between cooling and warming capacity

Cold-or warm-acclimation did not come at the expense of reduced heat or cold tolerance, respectively (Fig. 3). Specifically, there was no correlation between ECC and thermogenic capacity (i.e., *M*_sum_) in any treatment or at any age (3-4 weeks [cold: *t*_18_ = 0.10, p = 0.925, *r* = 0.02; warm: *t*_18_ = −1.06, p = 0.302, *r* = −0.24]; 8-9 weeks [cold: *t*_15_ = −0.91, p = 0.378, *r* = −0.23; warm: *t*_15_ = −0.41, p = 0.690, *r* = −1.05]; 12-13 weeks [cold: *t*_8_ = 1.74, p = 0.121, *r* = 0.52; warm: *t*_9_ = −0.69, p = 0.506, *r* = −0.23]).

**Figure 3.**
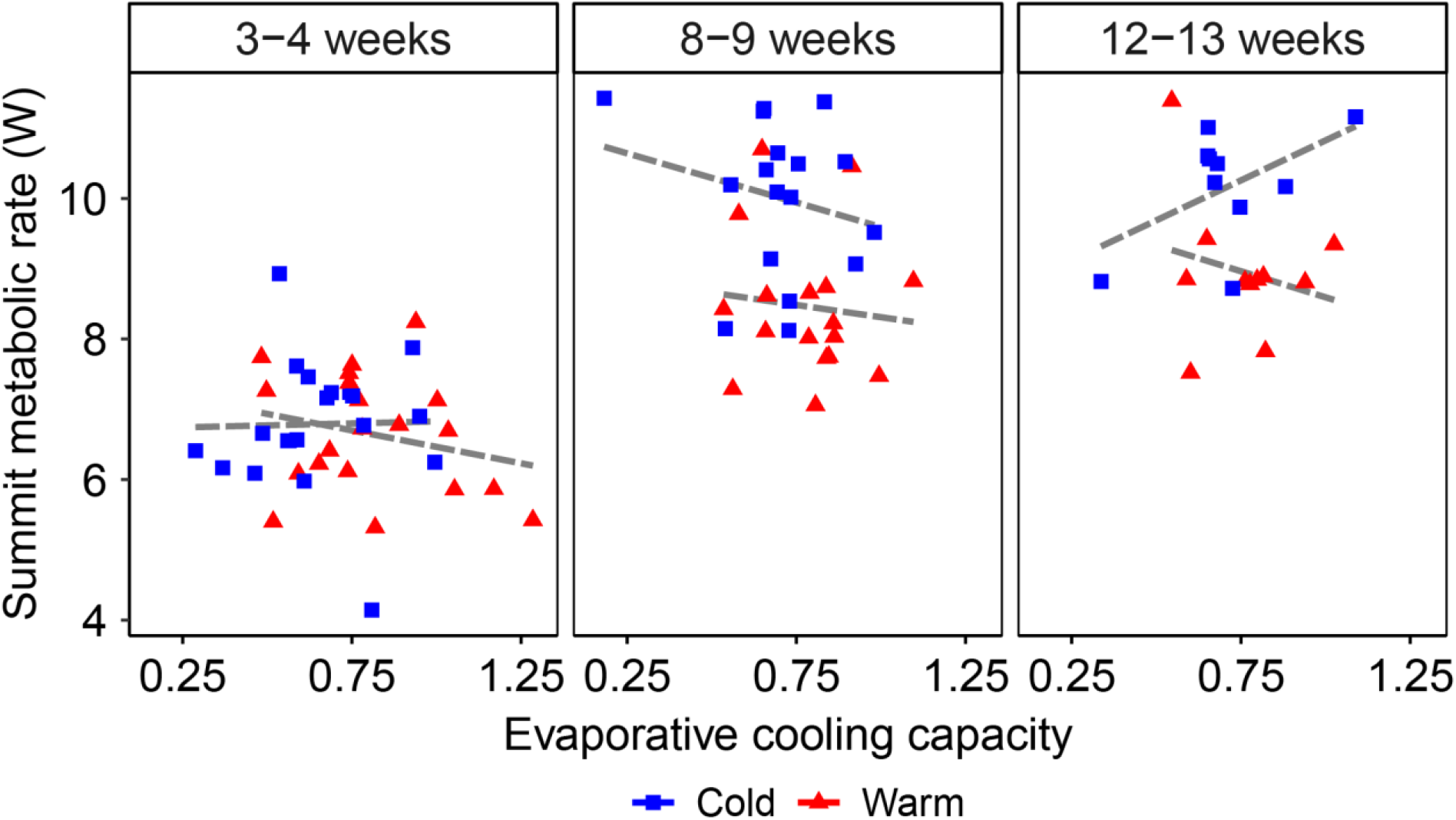
Relationship between maximum cold and heat tolerance in warm- and cold-acclimated Japanese quail. Cold tolerance was inferred from maximum thermogenic capacity (summit metabolic rate) during sliding cold exposure, and heat tolerance was inferred from the maximum evaporative cooling capacity during heat exposure. Lines represent correlations for each treatment and age (all P > 0.1). Sample sizes per age and temperature are stated in Table S1.

## Discussion

In line with our predictions, we found that Japanese quail growing up under cold snap-like conditions were more thermogenic and had improved cold tolerance limits compared to birds raised in the warmth. Development in heatwave-like conditions equipped birds with higher EWL and improved heat tolerance limits. While heat tolerance limits are typically far beyond temperatures birds are likely to experience over sustained periods in nature, it is plausible that individuals tolerating higher, or lower, temperatures can endure submaximal thermal stress with a lower somatic cost (cf. Orsted et al., 2022). This could allow the birds to maintain foraging effort, or exploit thermally challenging microhabitats, before having to actively engage in heat loss or heat gain processes, thus reducing missed-opportunity-costs of temperature variability (Cunningham et al., 2021).

Differences between cold snap- and heatwave-reared birds were largely reversed once the thermal stressor was removed and the birds had spent some weeks in common garden conditions. This might suggest that costs of reversible plasticity are less pronounced compared to in some ectotherm models (Morgan et al., 2022). The timeline of reversal differed between thermoregulatory traits (see Figs. 1, 2, S4), which might influence the capacity to deal with thermally labile environments. Moreover, the different timelines between trait reversals highlight the sometimes complex relationships between thermoregulatory traits and temperature tolerance limits (cf. Milbergue et al., 2018) and support recent work of the latter (Rego et al., 2025). Yet, previous research shows that birds can quickly acclimate both muscle ultrastructure and thermoregulatory performance when environmental temperature changes (Petit et al., 2013; Vezina et al., 2020). Further, reversal of temperature-dependent phenotypes aligns with previous endotherm studies showing that postnatal temperature conditions do not permanently program thermoregulation (Liew et al., 2003; Persson et al., 2024). It is likely that adaptive developmental priming requires close mapping in environmental temperature in juvenile and adult life stages. This may not be reasonable to assume for highly mobile animals like birds, where plasticity instead could ensure matching of the thermoregulatory phenotype to the surrounding thermal environment over short time frames. Alternatively, the endothermic nature of birds renders them physiologically or functionally homeothermic from an early age (e.g., Andreasson et al., 2016; Engert et al., 2025), a point beyond which any window for permanent programming may close. In line with this, the poikilothermic embryonic period appears more malleable to long-lasting developmental programming of temperature tolerance in birds (reviewed by Arjona et al., 1988; Piestun et al., 2008).

An outstanding question is whether heat- and cold tolerance are subjected to a trade-off, such that an individual can be either more thermogenic or more thermolytic, but not both. Likewise, there has been a recent call for understanding how any such trade-off may differ during development in line with ontogenetic changes in an individuals’ thermal relations (Schou & Cornwallis, 2024). Existing evidence for a trade-off is available at the level of reproductive investment, where ostriches that maintained high egg laying rate in the cold reduced investment in the warmth, and vice versa. The physiological basis for this relationship remains unstudied (Schou et al., 2022). While treatment groups differed expectedly in heat and cold responses, we found no correlation between heating and cooling capacity. Hence, while populations may vary in heat- and cold tolerance, we found no evidence for a trade-off in quail, possibly because thermogenesis and thermolysis follow largely distinct physiological pathways. Accordingly, a trade-off between heat- and cold tolerance may be apparent in some, but not, all traits. We advise this question is addressed further in a wider range of species across latitudes and biomes.

Thermoregulatory traits mediating cold tolerance increased with age in a treatment-specific manner, such that there were no significant differences in thermogenic capacity or cold tolerance limits in the juveniles (Fig. 1). This may reflect a trade-off between thermoregulation and growth. At the time of the first measurement (i.e., at 3 weeks), quail are in the middle of the exponential growth phase and roughly halfway to asymptotic size (Fig. S7). It is reasonable that the lower absolute amount of thermogenic tissue at this stage renders the birds less cold tolerant. Moreover, the juvenile plumage is often thinner and less insulating (Nord et al., 2023), which drives up heat loss rate (Nord & Folkow, 2018). When combining these predicaments, juvenile assimilation efficiency may simply be insufficient to maximise growth rate (required to reduce predation risk; e.g., Cheng & Martin, 2012; Martin, 1995) whilst simultaneously increasing investment in thermogenic tissue (e.g., Krijgsveld et al., 2003), explaining the lack of differences between treatments. At and beyond reproductive maturity, cold snap-reared birds displayed the expected increase in thermogenic performance and cold tolerance limit. It is noteworthy that this might not reflect increased mass of heat-producing tissue in Cold birds, since there was no overall difference in body size between the treatments (Figs. S6-S7; Tables S5-S6). Thus, our study supports the conclusion by Milbergue et al. (2018) that while larger muscles may be beneficial for heat production, it is not a strict necessity for improved cold tolerance. Future studies should investigate both the aerobic activity of skeletal muscle in cold-reared birds (cf. Liknes & Swanson, 2011) and any temperature-related differences in plumage insulation (cf. Osváth et al., 2018; Pap et al., 2020).

We have shown that, while postnatal developmental temperature impacts the ontogeny of thermoregulation, these effects appear to be plastic in entity. Moreover, and importantly, the scope for acclimating to cold and warm conditions differed. While the difference in cold tolerance limit between treatments was approximately 7°C, the increase in heat tolerance limit in Warm birds was only around 2°C. This was also apparent when comparing the common garden birds to their original thermal groupings: EWL and the heat tolerance limit was downregulated in Cold birds, but it was not *upregulated* in Warm birds. Similar results were recently found at the cellular level, where the thermal sensitivity of blood cell mitochondria were amenable to cold-acclimation but insensitive to warm-acclimation (Correia et al., 2025). While we can only speculate about the physiological processes mediating differences in acclimation capacity to heat and cold, it is likely that there is a hard ceiling for water turnover rate that precludes close matching of warming and ECC and makes birds sensitive to the detrimental effects of hyperthermia (reviewed by Schulte, 2015). As for ectotherms (Jorgensen et al., 2022; Morgan et al., 2020), this small scope for improving heat tolerance is worrying in a warming world. This may leave birds reliant on behavioural adjustment when physiology is exempted, with potentially severe downstream fitness consequences (cf. Cunningham et al., 2021).

## Supporting information

Electronic Supplementary Material

## ETHICS

Ethical approval was granted by the Malmö/Lund Animal Ethics Committee (acting under the authority of the Swedish Board of Agriculture; permit no. 9426-19).

## ACKNOWLEDGEMENTS

We are grateful for the comments improving previous versions of this article provided by Lars Råberg and Joshua K. R. Tabh. We thank Elisa Thoral, Joshua K. R. Tabh, Camilla Björklöv Andersson and Agniezska Czopek for help with bird care, and Lars Fredriksson for technical assistance.

## LIST OF ABBREVIATIONS

ECC: evaporative cooling capacity, defined as the ratio between evaporative heat loss and metabolic heat production
EHL: evaporative heat loss
ESM: electronic supplementary material
EWL: evaporative water loss
Helox: gas mixture of helium (79%) and oxygen (21%)
HTL: heat tolerance limit
MHP: metabolic heat production
*M*_sum_: summit metabolic rate
PIT: passive integrated transponder
*T*_b_: ambient temperature in metabolic chamber
*T*_b_: body temperature

## Additional Declarations

There is NO Competing Interest.

