## Supplementary material for "Postnatal developmental temperature affects the ontogeny of heat- and cold tolerance": Electronic Supplementary Material

### Experimental design

After an incubation period of 19 days at 37.5°C, 55 Japanese quail chicks were banded with uniquely numbered leg bands and were randomly allocated to either a cold snap-like (10°C) or a heatwave-like (30°C) housing regime (Table S1). The birds remained in these thermal environments until 9 weeks old. Then, half of the of the birds from each temperature group were transferred to a common garden at 20°C (henceforth Cold-mild or Warm-mild), whereas the other half remained in their original temperature treatments until the end of the experiment (Fig. S1; Table S1) to investigate if developmental temperature-effects reflected permanent programming or reversible plasticity. Body mass and wing length were measured throughout the experiment. Final sample sizes per treatment group, sex, and trait, are presented in Table S1.

The experimental design is illustrated in Fig. S1. We investigated how developmental temperature affected both cold- and heat tolerance throughout the experiment using flow-through respirometry (see main text for details). Cold tolerance was measured during sliding cold exposure (20°C per hour) using a helium-oxygen gas mixture (79% helium, 21% oxygen; helox) at 3, 8 and 12 weeks of age (see Fig. S2A for representative data). Heat tolerance was measured at 4, 9 and 13 weeks of age during incremental heat exposure (see Fig. S2B for representative data).

**Table S1.** The number of Japanese quail used for investigating morphological and physiological responses of developing in either Warm (30°C) or Cold (10°C) until reproductive maturity after which half of each treatment group were transferred into common garden conditions (Cold-mild and Warm-mild; 20°C).

| Age of birds (weeks) | Total number of birds | Treatment sample size | | | | Number of females/males | Sample size in analyses | | | | |
|---|---|---|---|---|---|---|---|---|---|---|---|
| | | Warm | Cold | Warm -mild | Cold -mild | | Body mass | Wing length | Daytime $T_b$ | Cold tolerance | Heat tolerance |
| 1 | 49 | 24 | 25 | - | - | 27 \| 22 | 49 | 49 | - | - | - |
| 2 | 49 | 24 | 25 | - | - | 27 \| 22 | 49 | 49 | - | - | - |
| 3 | 49 | 24 | 25 | - | - | 27 \| 22 | 49 | 49 | - | 47 | - |
| 4 | 48 | 23 | 25 | - | - | 26 \| 22 | 48 | - | - | - | 41 |
| 5 | 48 | 23 | 25 | - | - | 26 \| 22 | 48 | - | - | - | - |
| 6 | 48 | 23 | 25 | - | - | 26 \| 22 | 48 | - | 48 | - | - |
| 7 | 47 | 23 | 24 | - | - | 25 \| 22 | 47 | - | - | - | - |
| 8 | 47 | 23 | 24 | - | - | 25 \| 22 | 47 | 47 | - | 36 | - |
| 9 | 45 | 22 | 23 | - | - | 24 \| 21 | 45 | - | - | - | 41 |
| 10 | 44 | 11 | 10 | 11 | 12 | 24 \| 20 | 44 | - | - | - | - |
| 11 | 44 | 11 | 10 | 11 | 12 | 24 \| 20 | 44 | - | 42 | - | - |
| 12 | 44 | 11 | 10 | 11 | 12 | 24 \| 20 | 44 | 43 | - | 44 | - |
| 13 | 44 | 11 | 10 | 11 | 12 | 24 \| 20 | - | - | - | - | 43 |

Except for some experiments and measurements where birds were excluded, for reasons stated in the main text, the same birds were used at all ages and at all responses. Abbreviations: $T_b$: Body temperature.

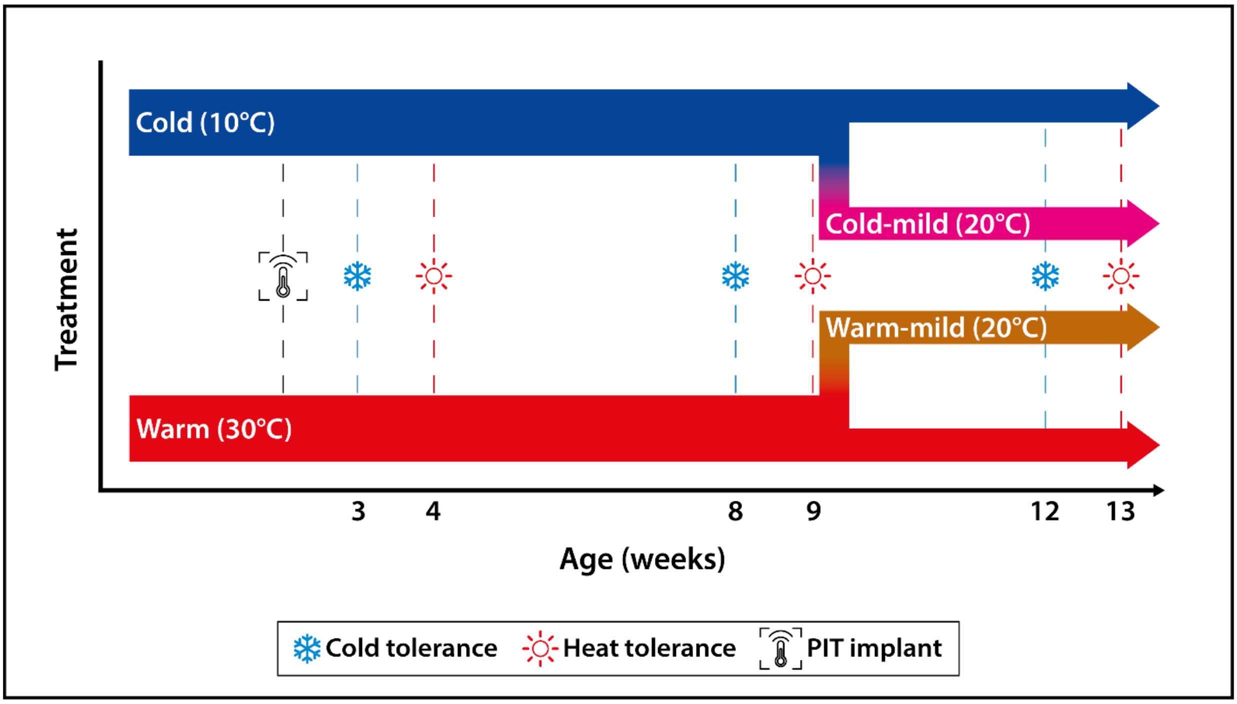

**Figure S1. Schematic overview of the experimental protocol.** Japanese quail were housed in either Warm (30°C) or Cold (10°C) conditions until 9 weeks of age, after which half of each treatment group were transferred to common garden conditions (Cold-mild and Warm-mild respectively; 20°C). Cold tolerance was measured at 3, 8 and 12 weeks of age and heat tolerance was measured at 4, 9 and 13 weeks of age. At 2 weeks of age, the birds were implanted with a passive integrated transponder (PIT) into the intraperitoneal cavity to measure body temperature, as described in the main text. Two days before each cold tolerance measurements, blood samples (< 1 % of blood volume) were collected from the brachial vein for another study.

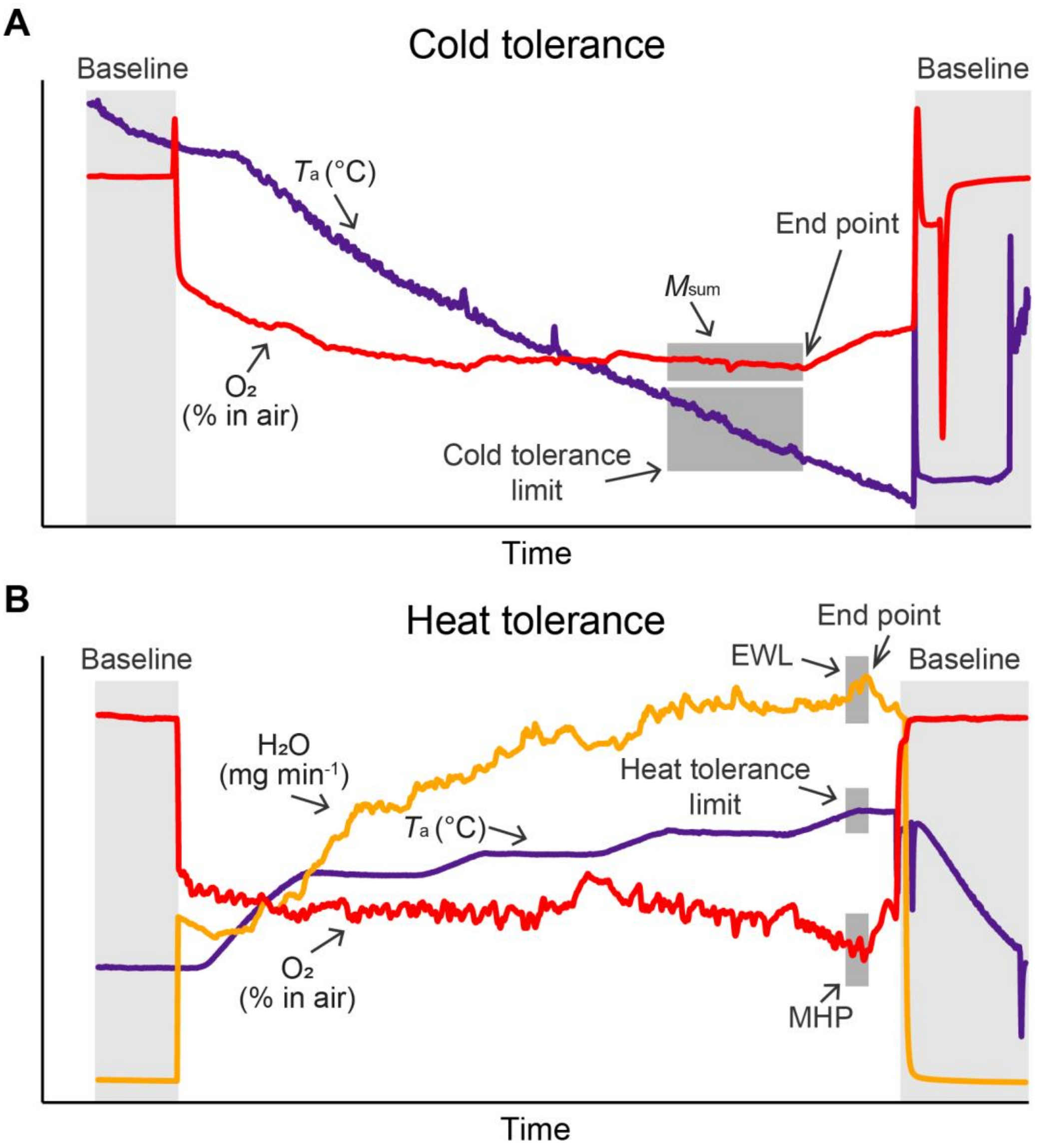

**Figure S2. Representative examples of (A) cold- and (B) heat tolerance measurements.** (A) A cold-tolerance experiment was performed in a helium-oxygen gas mixture (79% helium, 21% oxygen [$O_2$]). Ambient temperature in the climate chamber decreased by 20°C per hour. A bird was removed from the experiment (endpoint) when oxygen consumption started to decrease, or when there was no change in oxygen consumption for at least 20 min with decreasing ambient temperature ($T_a$). $O_2$-curve represents %$O_2$ in air, with increasing oxygen consumption %$O_2$ in air decreases while at the endpoint when $O_2$ consumptions starts to decrease %$O_2$ in air ($O_2$-curve) increases. The mean of the 10 min before the end point was used to calculate cold tolerance limit and $M_{sum}$ (shaded dark grey areas). (B) In heat tolerance measurements, after acclimation in 30°C (not depicted in the graph) $T_a$ was acutely increased to 40°C, after which $T_a$ increased in 2°C increments. A bird was assumed to have reached its' end point when oxygen consumption or water vapor concentration ($H_2O$; evaporative water loss) started to decrease, if body temperature ($T_b$) increased > 45°C, or if birds showed loss of coordination or signs of stress. The mean of the most level 2 min at stable $T_a$ before the endpoint was used for calculating the heat tolerance limit and its associated evaporative water loss and metabolic heat production (shaded dark grey areas). The shaded light grey areas show baseline measurements.

### Relationship between evaporative cooling capacity and heat tolerance limit

To investigate if evaporative cooling capacity (ECC) could be used to predict the heat tolerance limit (HTL), we analysed the relationship between ECC and HTL using linear regression (lm() in the stats package) with HTL as the dependent variable and ECC as the independent variable.

ECC significantly predicted HTL, which increased by 6.97°C at 4 weeks, 10.23°C at 9 weeks and 10.82°C at 13 weeks for each unit increase in the capacity for evaporative cooling (Fig. S3; Table S2).

**Table S2.** Parameter estimates from linear regressions between heat tolerance limit (i.e., the temperature above which a bird no longer could increase its evaporative water loss) and evaporative cooling capacity (ECC; i.e., the ratio between evaporative heat loss and metabolic heat production) in Japanese quail at 3 different ages. Estimates, test statistics, degrees of freedom and P-values at 4, 9 and 13 weeks of age.

| Model | Intercept | Slope | *t* | $R^2$ | Df | P |
|---|---|---|---|---|---|---|
| **Heat tolerance limit (°C) vs. ECC** | | | | | | |
| 4 weeks | 39.08 ± 1.005 | 6.97 ± 1.329 | 5.25 | 0.40 | 1,39 | <0.0001 *** |
| 9 weeks | 36.93 ± 0.850 | 10.23 ± 1.132 | 9.03 | 0.67 | 1,39 | <0.0001 *** |
| 13 weeks | 36.81 ± 1.570 | 10.82 ± 2.070 | 5.23 | 0.39 | 1,41 | <0.0001 *** |

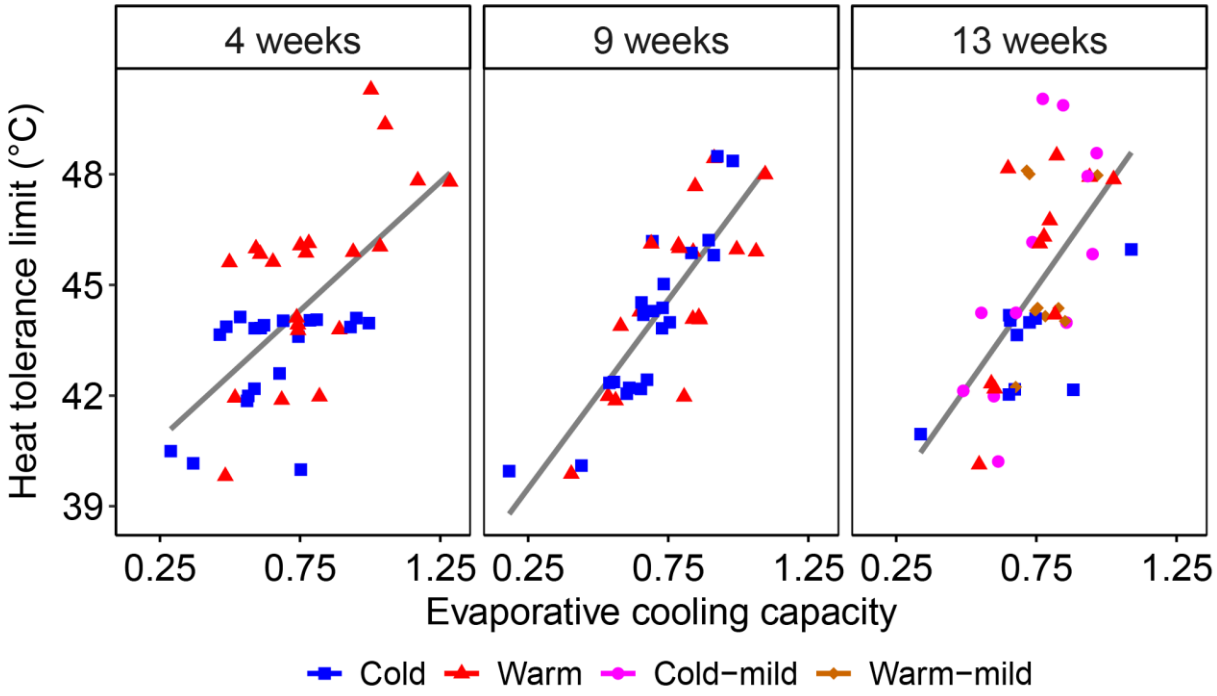

**Figure S3. Heat tolerance limit as a function of maximum evaporative cooling capacity at 4, 9, and 13 weeks of age in Japanese quail.** The birds were raised in either Warm (30°C) or Cold (10°C) conditions until 9 weeks of age, after which half of the quail were transferred to common garden conditions (Cold-mild, Warm-mild; 20°C). The other half remained in their original temperature treatments. Grey line shows regression line for respective age. Statistics are presented in Table S2. Sample sizes per age and temperature are stated in Table S1.

### Body temperature

Body temperature ($T_b$) was measured using a passive integrated transponder implanted into the intraperitoneal cavity (at 2 weeks of age; Fig. S1). This allowed readings of $T_b$ during tolerance measurements as well as daytime measurements in the holding pens. Daytime $T_b$ (08:00 – 20:00 GMT+2, i.e., 1 h after and before lights off) was measured twice, in-between experimental periods: firstly when the birds were 5-7 weeks old (mean ± s.e.m.: 43.7 ± 4.6 d), and secondly when the birds were 10-12 weeks old (77.2 ± 4.5 d). During these periods, birds were not handled and were left undisturbed apart from daily maintenance. Twelve to 2036 observations were collected per individual at each sampling point (mean: 489 ± 323 observations).

$T_b$ during tolerance measurements were analysed using linear mixed models with treatment, age, and treatment×age as factors. Effects of transfer to common garden was assessed using ANOVA comparing common garden birds with the respective origin treatment. Daytime $T_b$ was analysed using linear mixed models with treatment as a factor, mean centred body mass (by treatment, age and sex) as a covariate, and bird ID as random intercept. Separate models were fitted for the 5-7-week and 10-12-week periods. In the latter case, we compared Cold birds with Warm birds, Warm-mild birds with Warm birds and Cold-mild birds with Cold birds.

Daytime $T_b$ was higher in Warm compared to Cold birds (by 0.2 °C) in the first measurement period (i.e., at 5 to 7 weeks old; $p=0.010$; Fig. 4; Table S3), but this difference had disappeared by the second measurement period (at 10 to 12 weeks old). Nor was there a difference in daytime $T_b$ between the Warm-mild and Warm birds, and Cold-mild and Cold birds, respectively, at this age (Fig. S5; Table S3).

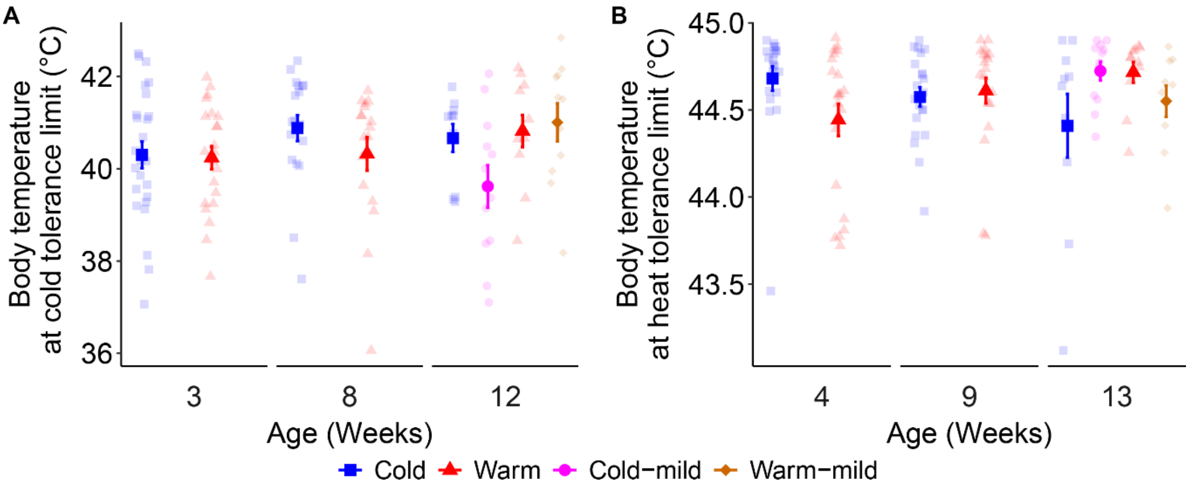

**Figure S4. Body temperature at (A) cold- and (B) heat tolerance limit at 3 different ages.** Cold tolerance was measured at 3, 8, and 12 weeks of age in 79% helium and 21% oxygen (helox). Heat tolerance was measured at 4, 9 and 13 weeks of age. Japanese quail developed in either Warm (30°C) or Cold (10°C) conditions until 9 weeks of age. Then, half of the quail were transferred to common garden (Cold-mild, Warm-mild; 20°C) and half remained in their origin temperature treatment. Semi-transparent points show raw data. Sample sizes per age and temperature are stated in Table S1.

**Table S3.** Parameter estimates explaining the effects on daytime $T_b$ of developing in either Warm (30°C) or Cold (10°C) temperature conditions in Japanese quail, after which half of the birds were transferred to common garden (Cold-mild, Warm-mild; 20°C) and the half remained in their origin treatment. Estimates, test statistics, degrees of freedom, P-values and standard deviations (σ) for the random factor from linear mixed models at 5-7 and 10-12 weeks on daytime $T_b$. Different letters within brackets represent significant (P < 0.05) post hoc comparisons. Abbreviations: $T_b$: Body temperature.

| **Model** | **Estimate ± s.e.m** | **LR** | **Df** | **P** | **$\sigma_{ID}$\|$\sigma_{total}$** |
|---|---|---|---|---|---|
| **Daytime $T_b$** | | | | | |
| 5-7 weeks | | | | | |
| Treatment | | 6.57 | 1 | 0.0104 * | |
| Cold [A] | 41.97 ± 0.059 | | | | |
| Warm [B] | 42.19 ± 0.061 | | | | |
| Body mass | -0.005 ± 0.002 | 6.24 | 1 | 0.0125 * | |
| Bird ID (random) | | 15361.00 | 1 | <0.0001 *** | 0.06\|0.15 |
| 10-12 weeks Cold~Warm | | | | | |
| Treatment | | 0.60 | 1 | 0.4347 | |
| Body mass | 0.005 ± 0.002 | 8.71 | 1 | 0.0032 * | |
| Bird ID (random) | | 4457.30 | 1 | <0.0001 *** | 0.06\|0.31 |
| 10-12 weeks Cold~Cold-mild | | | | | |
| Treatment | | 1.56 | 1 | 0.2113 | |
| Body mass | | 0.05 | 1 | 0.8259 | |
| Bird ID (random) | | 10493.00 | 1 | <0.0001 *** | 0.07\|0.34 |
| 10-12 weeks Warm~Warm-mild | | | | | |
| Treatment | | 3.24 | 1 | 0.0717 | |
| Body mass | | 3.03 | 1 | 0.0818 | |
| Bird ID (random) | | 9768.60 | 1 | <0.0001 *** | 0.09\|0.40 |

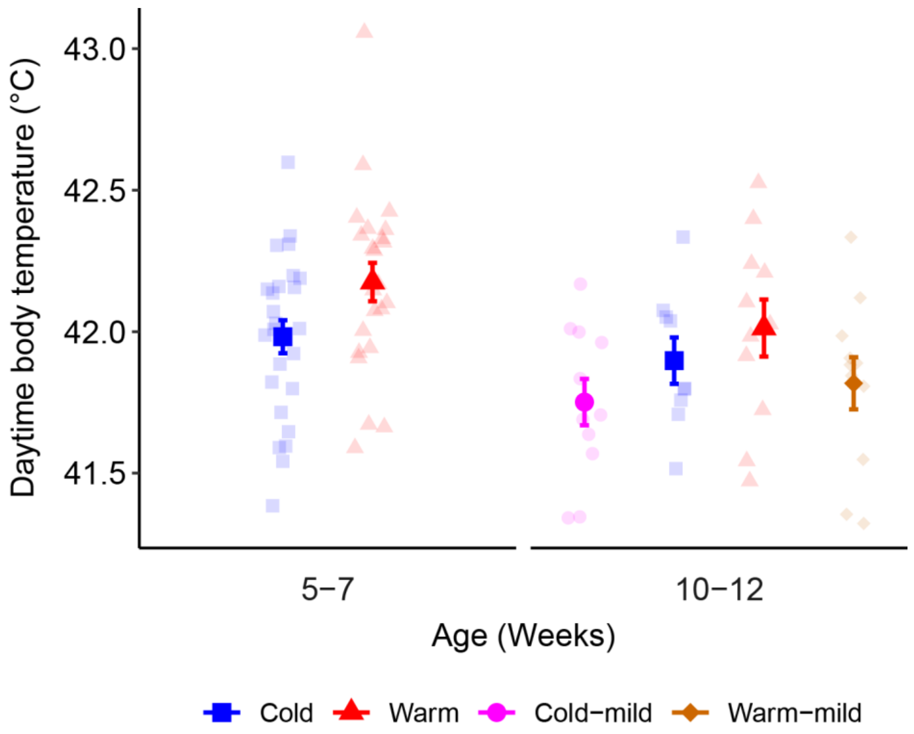

**Figure S5. Body temperature at daytime between tolerance measurements periods (mean ± s.e.m.).** Japanese quail developed in either Warm (30°C) or Cold (10°C) conditions until 9 weeks of age, then half of the quail moved to common garden (Cold-mild, Warm-mild; 20°C) and half remained in their origin temperature treatment. Semi-transparent points show raw data. Sample sizes per age and temperature are stated in Table S1.

### Body mass and wing length

Body mass (± 0.1 g) and wing length (± 0.5 mm) were measured once weekly and at 1, 2, 3, 8, and 12 weeks of age, respectively, to investigate any effects of the developmental temperature treatment on somatic growth and size. To test if body mass and wing length differed between treatment groups and sex at reproductive maturity (9 and 8 weeks, respectively), and at 12 weeks of age, ANOVAs (anova() function in stats package) were used, with treatment, sex, and treatment×sex as factors. To test if there was an effect of transfer to common garden conditions, Cold-mild and Warm-mild birds were teste against their origin treatment.

Next, to analyse if growth was affected by treatment, we fitted linear mixed models (lmer() in lme4; Bates et al., 2015) with either body mass or wing length as dependent variables, treatment as a factor, and age and $age^2$ as covariates. The interactions treatment×age and treatment×$age^2$ were included in the in the first models but was removed since they were non-significant ($p>0.05$). Bird ID was used as random factor to account for repeated measurements.

Cold- and heat-acclimation affected body mass at 9 weeks in a sex-specific manner, where sexual size dimorphism was less pronounced in Warm compared to Cold birds (Fig. S6A; Table S4). Specifically, Cold males were 22% lighter than Cold females, but Warm males only tended to be lighter than Warm females (by 11%; $p = 0.069$; Fig. S6A; Table S4). Also, Cold males were lighter than Warm females and Warm males were lighter than Cold females (20% and 16% respectively; Fig. S6A; Table S4). However, there were no significant difference between Warm and Cold females. At 12 weeks of age, females weighed more than males across groups, but there was no effect of treatment on body mass (Figs. S6C, S6E; Table S4). Nor was growth rate over the 12-week experiment affected by cold- and warm-acclimation (Fig. S7A, Table S5).

At 8 weeks of age (at adult size [Fig. S7B; Table S5]) there was no effect of treatment on wing length, but males had significantly shorter wings than females (by 2%; Fig. S6B; Table S4). Warm birds had 4% longer wings than Cold birds at 12 weeks and, again, females had longer wings than males (by 4%; Fig S6D; Table S4). Wing length did not differ between Cold and Cold-mild birds at 12 weeks. When comparing Warm and Warm-mild birds, Warm males had shorter 4% wings than Warm females, but there was no effect of sex in the Warm-mild group (treatment×sex, $p = 0.017$). Growth rate over the 12-week experiment was affected by treatment, with Warm birds having longer wings than cold birds (Fig. S7B, Table S5).

**Table S4.** Parameter estimates explaining the effects of developing in either Warm (30°C) or Cold (10°C) temperature conditions until 9 weeks on body mass and wing length in Japanese quail. After which half of each treatment group were transferred to common garden (Cold-mild, Warm-mild; 20°C), and the other half remained in their original treatment. Estimates, test statistics, degrees of freedom and P-values from ANOVAs on body mass and wing length. Significant (P < 0.05) post hoc comparisons are shown by different letters within brackets.

| Model | Estimate ± s.e.m. | F | Df | P |
|---|---|---|---|---|
| **Body mass (g)** | | | | |
| Cold~Warm – 9 weeks | | | | |
| Treatment:Sex | | 8.27 | 1,41 | 0.0064 ** |
| Cold | | | | |
| Female [A] | 299.2 ± 8.10 | | | |
| Male [B] | 223.2 ± 11.21 | | | |
| Warm | | | | |
| Female [A] | 279.5 ± 11.00 | | | |
| Male [A] | 250.0 ± 16.16 | | | |
| Cold~Warm – 12 weeks | | | | |
| Treatment | | 0.99 | 1,19 | 0.3323 |
| Sex | | 19.90 | 1,19 | 0.0003 *** |
| Female [A] | 287.0 ± 8.04 | | | |
| Male [B] | 232.3 ± 12.28 | | | |
| Treatment:Sex | | 2.51 | 1,17 | 0.1309 |
| Cold~Cold-mild – 12 weeks | | | | |
| Treatment | | 1.39 | 1,20 | 0.2521 |
| Sex | | 31.01 | 1,20 | <0.0001 *** |
| Female [A] | 308.7 ± 10.09 | | | |
| Male [B] | 229.2 ± 14.27 | | | |
| Treatment:Sex | | 0.21 | 1,18 | 0.6538 |
| Warm~Warm-mild – 12 weeks | | | | |
| Treatment | | 0.46 | 1,20 | 0.5038 |
| Sex | | 15.09 | 1,20 | 0.0009 *** |
| Female [A] | 292.9 ± 6.71 | | | |
| Male [B] | 252.1 ± 10.45 | | | |
| Treatment:Sex | | 0.47 | 1,18 | 0.5025 |
| **Wing length (mm)** | | | | |
| Cold~Warm – 8 weeks | | | | |
| Treatment | | 2.38 | 1,45 | 0.1298 |
| Sex | | 5.98 | 1,45 | 0.0184 * |
| Female [A] | 124.8 ± 0.68 | | | |
| Male [B] | 122.3 ± 0.99 | | | |
| Treatment:Sex | | 2.49 | 1,43 | 0.1222 |
| Cold~Warm – 12 weeks | | | | |
| Treatment | | 8.47 | 1,19 | 0.0090 ** |
| Cold [A] | 120.2 ± 1.14 | | | |
| Warm [B] | 124.8 ± 1.16 | | | |
| Sex | | 12.39 | 1,19 | 0.0023 ** |
| Female [A] | 124.8 ± 0.97 | | | |
| Male [B] | 119.6 ± 1.48 | | | |
| Treatment:Sex | | 0.05 | 1,17 | 0.8303 |
| Cold~Cold-mild – 12 weeks | | | | |
| Treatment | | 3.55 | 1,19 | 0.0749 . |
| Sex | | 10.55 | 1,19 | 0.0042 ** |
| Female [A] | 124.2 ± 1.03 | | | |
| Male [B] | 119.4 ± 1.49 | | | |
| Treatment:Sex | | 0.04 | 1,17 | 0.8416 |
| Warm~Warm-mild – 12 weeks | | | | |
| Treatment:Sex | | 6.96 | 1,18 | 0.0167 * |
| Warm | | | | |
| Female [A] | 126.7 ± 0.97 | | | |
| Male [B] | 121.6 ± 1.61 | | | |
| Warm-mild | | | | |
| Female [A] | 124.8 ± 1.43 | | | |
| Male [A] | 132.5 ± 2.24 | | | |

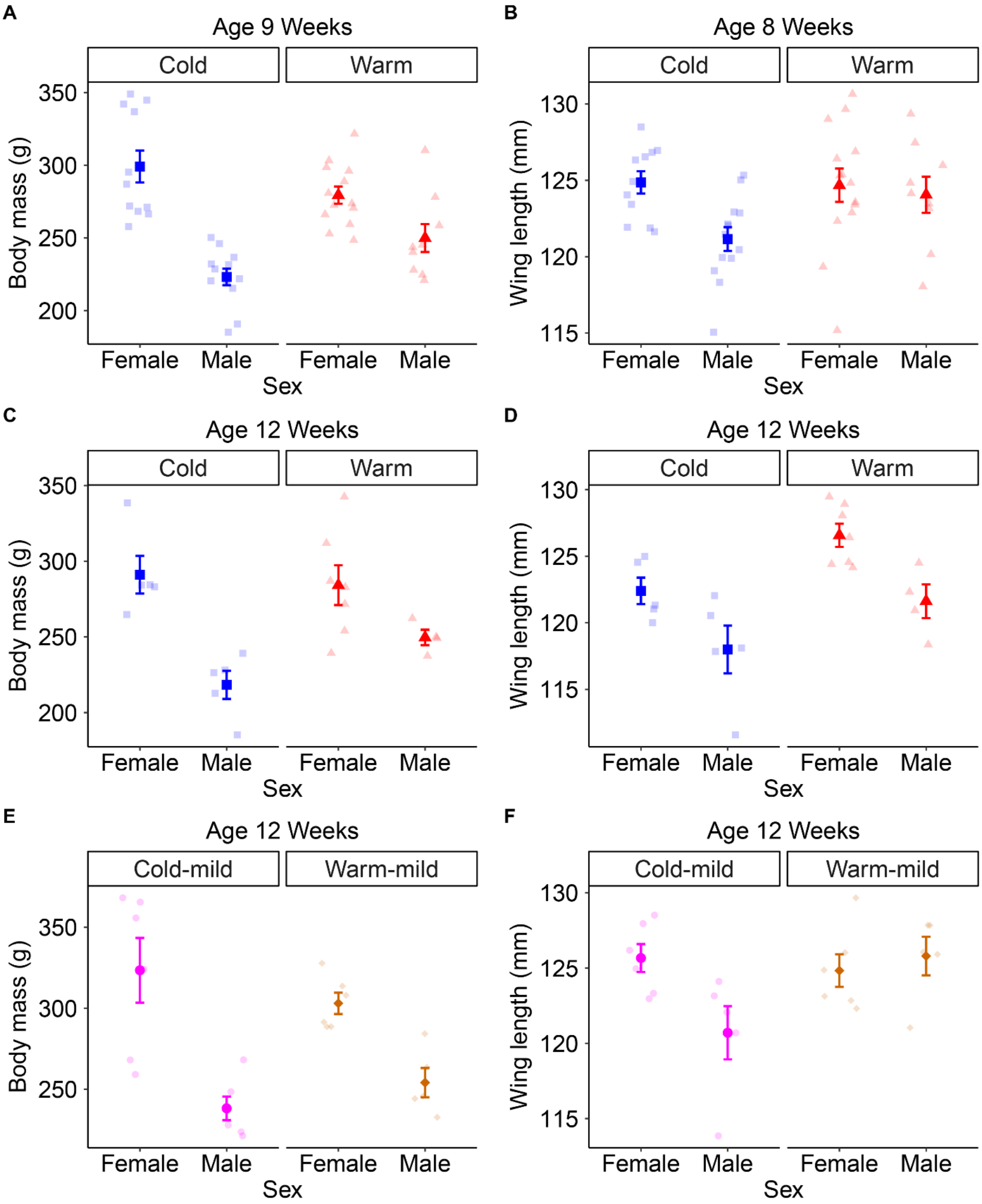

**Figure S6. (A) Body mass and wing length at 9 and 8, and 12 weeks of age (mean ± s.e.m.).** (A) Body mass at 9 weeks of age and (B) wing length at 8 weeks of age of Japanese quail that developed in either Warm (30°C) or Cold (10°C) conditions until 9 weeks of age. (C) Body mass and (D) wing length at 12 weeks of age of the half of quail that remained in their temperature treatment until end of experiment, and (E) body mass and (F) wing length at 12 weeks of age of the other half of quail that was transferred to common garden conditions (Cold-mild, Warm-mild; 20°C) until end of experiment. Semi-transparent points show raw data. Sample sizes per age and temperature are stated in Table S1.

**Table S5.** Parameter estimates explaining the effects of developing in either Warm (30°C) or Cold (10°C) temperature conditions on body mass and wing length in Japanese quail. Estimates, test statistics, degrees of freedom, P-values and standard deviations (σ) for the random factor from linear mixed models on body mass and wing length.

| **Model** | **Estimate ± s.e.m.** | **LR** | **Df** | **P** | $\sigma_{ID} \vert \sigma_{total}$ |
|---|---|---|---|---|---|
| Body mass (g) | | | | | |
| (Intercept) | -65.69 ± 4.825 | | | | |
| Treatment | | 1.52 | 1 | 0.2184 | |
| Age | 60.50 ± 0.917 | 1141.22 | 1 | <0.0001 *** | |
| Age$^2$ | -2.83 ± 0.145 | 723.17 | 1 | <0.0001 *** | |
| Treatment:Age | | 0.10 | 1 | 0.7523 | |
| Treatment:Age$^2$ | | 0.41 | 1 | 0.5208 | |
| Bird ID (random) | | 298.25 | 1 | <0.0001 *** | 19.82\|37.95 |
| Wing length (mm) | | | | | |
| (Intercept) | -2.85 ± 2.794 | | | | |
| Treatment | 4.38 ± 1.425 | 9.43 | 1 | 0.0021 ** | |
| Age | 28.26 ± 1.020 | 329.91 | 1 | <0.0001 *** | |
| Age$^2$ | -1.49 ± 0.071 | 241.44 | 1 | <0.0001 *** | |
| Treatment:Age | | 0.93 | 1 | 0.3342 | |
| Treatment:Age$^2$ | | 0.77 | 1 | 0.3807 | |
| Bird ID (random) | | 0.00 | 1 | 1.0000 | 0.00\|0.00 |

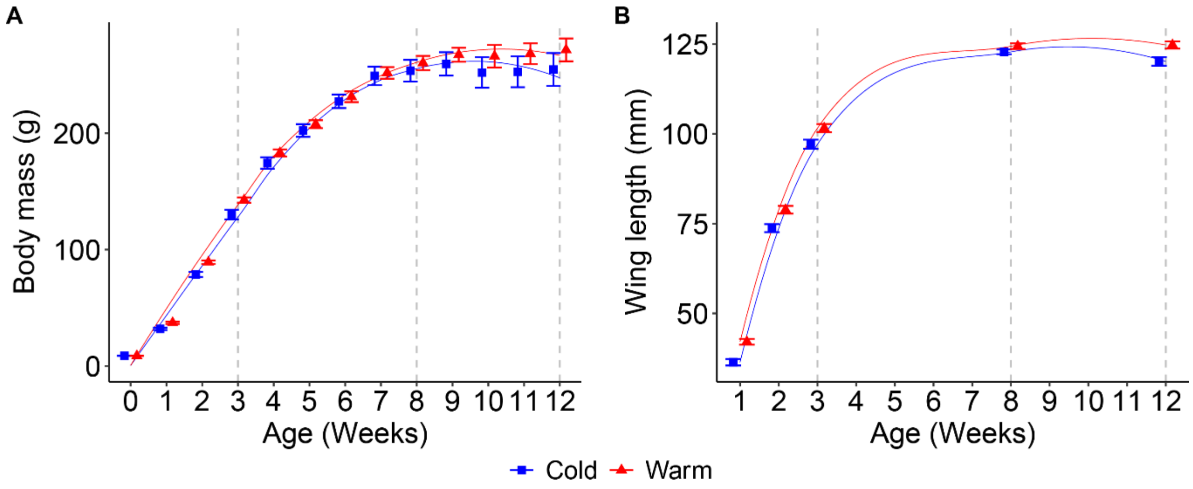

**Figure S7. (A) Body mass and (B) wing length at 1 to 12 weeks of age (mean ± s.e.m.).** Japanese quail developed in either Warm (30°C) or Cold (10°C) conditions until 9 weeks of age, then half of the quail moved to common garden (Cold-mild, Warm-mild; 20°C) and half remained in their origin temperature treatment. The curves are LOESS (locally estimated scatterplot smoothing). The dashed lines represent the timing of cold tolerance measurements. Sample sizes per age and temperature are stated in Table S1.
